# PreS1-decorated recombinant adenovirus encoding HBV antigens generates neutralizing humoral and cellular immunity

**DOI:** 10.64898/2026.01.30.702816

**Authors:** Rebecca A. Russell, James Lok, James M. Harris, Senko Tsukuda, Louisa M. Rose, Bethany G. Charlton, Ivana Carey, Kosh Agarwal, Peter A.C. Wing, Sumi Biswas, Jane A. McKeating, Matthew D. J. Dicks

## Abstract

**Background & Aims:** Achieving functional cure for chronic hepatitis B (CHB) will likely require a combinatorial approach targeting multiple aspects of the complex hepatitis B virus (HBV) life cycle and host adaptive immune responses. Here, we developed a therapeutic vaccination strategy targeting the PreS1 region of L-HBsAg, required for cellular entry of both hepatitis B and D viruses. An established potent T-cell inducing platform, recombinant adenovirus (Ad), was used as a nanoparticle scaffold for PreS1 attachment, to generate antibodies that neutralize virus entry and to establish T-cell mediated immune control.

**Approach & Results:** Screening a cohort of 61 patients diagnosed with CHB revealed minimal evidence of natural anti-PreS1 responses. Thus, Ad particles encoding multiple HBV antigens were decorated with PreS1 peptide using DogTag/DogCatcher protein superglue. Mice vaccinated with PreS1-decorated Ad induced robust anti-PreS1 antibody responses that neutralized HBV and HDV infection. In contrast, an undecorated Ad encoding L-HBsAg failed to neutralize HBV, demonstrating that PreS1 decoration was required for potent HBV neutralization. Strong CD8+ and CD4+ T-cell responses were induced against HBV antigens encoded in the Ad genome.

**Conclusions:** PreS1-decorated Ad combines immunological HBV and HDV entry inhibition with potent anti-HBV T-cell induction in a single platform, providing a promising addition to current therapeutic strategies against CHB, with particular utility in HBV/HDV co-infection.

## Introduction

In 2022, the WHO estimated that approximately 254 million people worldwide were living with chronic hepatitis B (CHB) infection, leading to approximately 1.1 million deaths annually from liver cirrhosis and cancer (1). Current treatments for CHB include nucleos(t)ide analogues (NA), and pegylated interferon alpha (2). These treatments are not curative. NA treatments suppress reverse transcription of HBV pregenomic RNA, but have no direct effect on covalently closed circular DNA; as a result, peripheral viral loads frequently rebound on NA cessation (3). Current prophylactic HBV vaccines, based primarily on the HBV small surface antigen (S-HBsAg), induce protective antibody responses in HBV naïve individuals that can prevent infection, but have limited efficacy against pre-existing chronic infection (4). One potential explanation is that in CHB, high levels of circulating S-HBsAg lead to antigen-specific immune tolerance (5) and exhaustion of S-HBsAg-specific B- and T-cells (6). There remains an unmet need for an effective treatment capable of inducing a functional cure for CHB, defined as sustained HBsAg loss and undetectable serum HBV DNA, with or without anti-HBs seroconversion (7).

The PreS1 region of the HBV large surface antigen (L-HBsAg) has been proposed as a target antigen for therapeutic HBV vaccination. PreS1 is essential for virus entry into target cells through interactions with the cellular receptor for HBV and HDV (sodium taurocholate co-transporting polypeptide (NTCP)) (8, 9). Antibodies against PreS1 can inhibit HBV and HDV infection (10-12). Furthermore, PreS1-specific B- and T-cells may be less functionally exhausted in CHB than those recognising S-HBsAg (13), since circulating levels of L-HBsAg tend to be far lower than S-HBsAg; where non-infectious subviral particles rich in S-HBsAg can exceed infectious Dane particles by several orders of magnitude during chronic infection (14). Earlier studies have suggested that anti-PreS1 antibody responses are typically weak or absent in CHB patients, thus presenting an opportunity for therapeutic intervention (15, 16). Indeed, PreS1 seroconversion has been proposed as an early prognostic marker for resolution of acute infection, rarely observed in patients progressing to chronic infection (16). Recently, Wang *et al* (17) used SpyTag and SpyCatcher protein superglue technology (18) to attach PreS1 to a ferritin nanoparticle. PreS1-displaying nanoparticles neutralized HBV infection *in vitro* and significantly reduced levels of HBsAg and HBV DNA in an adeno-associated virus-based murine model of HBV in both prophylactic and therapeutic settings (17).

A therapeutic strategy inducing neutralizing anti-PreS1 antibodies could limit the spread of infection, inhibit the replenishment of viral reservoirs (19) and may target infected cells through antibody-dependent cellular cytotoxicity (20). However, it is widely accepted that establishment of T-cell-mediated immune control will be important for successful therapeutic HBV vaccination (21). HBV-specific CD8+ T-cell responses are critical for resolution of acute infection (22) but are dysfunctional during chronic disease. Recombinant adenovirus (Ad) vectors are among the most potent inducers of T-cell immunity, particularly CD8+ T-cell responses, in humans (23, 24). Recently, a chimpanzee Ad encoding multiple HBV antigens induced robust viral-specific T-cell responses in healthy volunteers and lower but significant responses in CHB patients after a single administration (25). Combining this vaccine with a Modified Vaccinia virus Ankara (MVA) vector encoding the same HBV antigens in a prime-boost regimen generated stronger T-cell responses in CHB patients and significantly reduced surface antigen expression in a subset of patients with lower baseline HBsAg (26). Although humoral immunity was not measured in either of these clinical studies (25, 26), HBs antibody responses were modest in preclinical studies with these viral vector vaccines (27). Another promising approach currently in Phase I trials is TherVacB, a heterologous prime-boost vaccination strategy using particulate HBsAg and HBV core antigen (HBcAg) protein vaccines and an MVA vector encoding the same antigens. This regimen elicits strong humoral and cellular immunity and has been shown to reduce peripheral HBsAg and HBV DNA levels in HBV mouse models (28). Importantly, TherVacB does not contain PreS1 and HBV mouse models do not fully recapitulate chronic inflammatory disease in humans, since they often lack an adaptive immune system or do not support the complete replicative HBV life cycle.

Building on the approaches described above, we created a recombinant Ad vector encoding L-HBsAg and HBcAg and simultaneously decorated the Ad with PreS1 peptide. HBcAg forms the internal capsid structure of HBV particle, and HBcAg-specific T-cells have been shown to be more abundant and less functionally exhausted than S-HBsAg specific T-cells in CHB (29, 30). Covalent attachment of PreS1 to the Ad capsid surface was achieved using the DogTag/DogCatcher protein superglue system, which is similar to SpyTag/SpyCatcher but performs more efficiently in capsid loop structures (31). We previously introduced modular capsid decoration as a strategy to improve the potency and durability of humoral immune responses by attaching target antigens for pathogen neutralization to the Ad particle surface (32). Importantly, decorated Ad retained the ability to induce robust CD8+ T-cell immunity against encoded transgene antigens (32).

In the present study, immunization of mice with PreS1-decorated Ad induced superior serum responses that could neutralize HBV compared to an undecorated Ad encoding L-HBsAg. Decorated Ad particles induced comparable T-cell responses against encoded antigens to undecorated particles. Our therapeutic vaccine concept, combining PreS1-targeted neutralization of infectious HBV with induction of cellular immunity using a potent T-cell-inducing Ad-based platform has the potential to augment current therapeutic strategies against CHB.

## Materials and Methods

### Cloning and production of adenoviral vectors encoding HBV genes

Construction of Ad-DogTag, an E1/E3 deleted (replication defective) Ad serotype 5 vector with DogTag inserted into hexon hypervariable loop 5 (HVR5) was described previously (32). Ad-DogTag constructs expressing combinations of S-HBsAg, L-HBsAg and HBcAg, from genotype D, serotype ayw NC_003977 (human codon optimized) were generated through cloning of gene constructs into pENTR4.CMVp and subsequent Gateway-mediated insertion into the E1 locus using Invitrogen Gateway site-specific recombination technology. HBV gene cassettes are described in Figure S1. Recombinant Ad-DogTag vectors were produced, purified and titered as described previously (32).

### Protein production and purification

DogCatcher-PreS1 was expressed in suspension ExpiCHO-S cells (ThermoFisher, UK), affinity purified using C-tag affinity resin (ThermoFisher, UK) on an AKTA chromatography system (GE Healthcare, UK) and dialyzed into Tris-buffered saline pH 7.8. PreS1 without DogCatcher, cloned into pGEX-6P-1, was transformed into BL21DE3 cells (Life Technologies, UK). The GST-PreS1 protein was purified from the *E*.*coli* pellet using Glutathione Spin columns (Pierce, UK) following the manufacturer’s instructions. The PreS1 protein was then separated from GST by PreScission Protease (GE Healthcare, UK) following the manufacturer’s on-column cleavage protocol. Purified PreS1 was dialyzed into Tris-buffered saline pH 7.8.

### Coupling reactions

For *in vitro* assays, coupling reactions between DogCatcher-PreS1 protein and Ad-DogTag were performed by co-incubation of 1E+10 viral particles with DogCatcher-PreS1 at a range of μM concentrations. Reactions were incubated for 16 h at 4 °C in a total volume of 20 μL sucrose storage buffer (10 mM Tris-HCl, 7.5% w/v sucrose, pH 7.8). Ligand-decorated vector batches for vaccine studies were prepared by scaling up the *in vitro* reactions from 20 μL to 1 mL. Excess ligand was removed using SpectraPor dialysis cassettes with a 1000-kDa molecular weight cutoff (Spectrum Labs, UK). Dialysis decreased the excess ligand by at least 10-fold, as measured by densitometry on Coomassie-stained SDS-PAGE. Ligand coverage was determined by SDS-PAGE as described previously (32).

### Mouse immunizations

All mouse procedures were performed in accordance with the terms of the UK Animals (Scientific Procedures) Act (Project License PP5949437) and approved by the Oxford University Ethical Review Body. Female C57BL/6 mice, 6 per group (aged 6–8 weeks, Envigo, UK), housed in specific pathogen-free environments, were immunized intramuscularly by injection of 50 μL of vaccine formulated in endotoxin-free PBS (Gibco, UK) into both hind limbs of each animal (100 μL in total). Ad vector vaccine doses administered were 1E+8 infectious units (ifu)/animal/immunisation. The DogCatcher-PreS1 protein vaccine dose was 0.2 μg/animal/immunisation, equivalent to the PreS1 antigen dose delivered by Ad:PreS1^high^ vaccination, based on the capsid antigen dose calculation described previously (32). Protein vaccines were formulated with Addavax (InvivoGen, USA) at a 1:1 v/v ratio of adjuvant to antigen. Endotoxin dose was <3 EU per mouse in all studies. Experiments were performed at the Wellcome Trust Centre for Human Genetics, Functional Genomic Facility, University of Oxford.

### IgG ELISA

IgG endpoint ELISA was performed as described previously (33) with plates coated with 0.5 μg/mL PreS1. Plates were blocked with 10% Milk powder in 0.05% PBS-Tween20. Serum was diluted 3-fold in 0.05% PBS-Tween20 with a starting dilution of 1 in 100. Secondary antibody (STAR117A, Bio-Rad, UK) was diluted 1 in 5000 and the signal was developed in *p*-nitrophenylphosphate solution. The signal was allowed to develop for 30 minutes and plates were read at 405 nm on an Infinite M Plex (Tecan, UK). Data were analysed in Prism v10.

### Ex vivo IFNγ-ELISpot

Overnight *ex vivo* IFNγ-ELISpot was performed on freshly isolated splenocytes according to standard protocols, as described previously (34). To measure antigen-specific responses, cells were re-stimulated with peptides at a final concentration of 5 mg/ml for 18–20h. To measure T-cell responses to GFP, CD8+ T-cell epitope peptide EGFP_118-126_ (35) was used. To measure CD8+ T-cell responses to L-HBsAg and S-HBsAg, cells were stimulated with Env_190-197_ (36). To measure CD8+ T-cell responses to HBcAg, cells were stimulated with Core_93-100_ (37). Spot-forming cells were measured using an automated ELISpot reader system (AID Classic, Germany) with software version 7.

### Splenocyte stimulation, intracellular cytokine staining and flow cytometry

Frozen mouse splenocytes were thawed and rested overnight. Next day the splenocytes were washed, counted, seeded at 1x10^7^ cells/ml in the presence of 1 μL/mL GolgiPlug (BD Biosciences, UK) and a specific stimulation peptide or peptide pool at 1 μg/mL for each peptide. To measure T-cell responses to HBV L, a pool of 95 15-mer peptides with an 11-mer overlap was used (ProImmune, UK). To measure T-cell responses to HBV Core, PepMix HBV (Capsid Protein) Ultra (JPT, Germany) consisting of 155 individual 15-mer peptides with an 11-mer overlap was used. As a positive control each splenocyte sample was incubated with 1 μL/mL GolgiPlug and 20 ng/mL phorbol 12-myristate 13-acetate and 1 μg/mL Ionomycin. After 6 hours of incubation at 37°C with 5% CO_2_ the cells were stored at 4°C overnight. Next day the cells were stained. See supplemental methods for staining protocol and antibody panel information. Samples were analysed on an Attune NxT Flow Cytometer (ThermoFisher, UK) and analysed in FlowJo v 10.10 (BD Biosciences, UK).

### HBV Neutralization assay

Heat-inactivated sera from mouse experiments were pooled and incubated for 1 hour at 37°C with an HBV reporter virus (multiplicity of infection - 600) expressing Gaussia Luciferase (rHBV-gLuc), before being added to HepG2-NTCP cells (9) (see supplemental methods for HepG2-NTCP cell preparation). Briefly, rHBV-gLuc is a heparin-purified (38), single cycle virus, using authentic HBV entry pathways, but is replication-incompetent as a Gaussia Luciferase gene has been engineered in place of the viral polymerase (39). Six days post infection the level of Gaussia Luciferase was measured by mixing 60 μL cell culture supernatant with an equal volume of coelenterazine substrate (500 μg/mL) (Promega, UK) in a 96-well white plate (Corning, USA) and immediately reading the plate on a FluoStar (BMG Labtech, UK) with 1 s integration.

### HDV neutralisation assay

Heat-inactivated sera from mouse experiments were pooled, diluted to 1:125 and incubated for 1 hour at 37°C with HDV (multiplicity of infection - 50 genome copies per cell) before being added to HepG2-NTCP cells. Six days post infection HDV-infected cells were detected with an anti-HDAg antibody (pan-genotypic antibody [FD5C4], Kerafast, USA) and nuclei were detected with DAPI. See supplementary methods for HDV preparation and HepG2-NTCP cell preparation.

### CHB virus patient sample preparation

All patients had provided written consent for blood samples to be collected by the King’s College Hospital Liver Research Biobank and used for research purposes (IRAS project 332608; REC reference 23/LO/0708). All research was conducted in accordance with both the Declarations of Helsinki and Istanbul. Collection tubes were inverted 6-8 times, and then centrifuged at 2200 × g for 15 min. The supernatant was stored at -20°C in the biobank repository.

### Statistics

Statistical analyses were performed in GraphPad Prism v10. Comparisons between multiple groups were performed by Kruskal-Wallis with Dunn’s test for multiple comparisons. Pair-wise comparisons (two groups only) were performed with the Mann Whitney test. Statistical significance is indicated as follows: *p < 0.05; **p < 0.01; ***p < 0.001; ****p < 0.0001; non-significant comparisons are not shown. Where only certain statistical comparisons have been made this is described in the figure legend.

## Results

### PreS1 antibody titers are rarely observed in patients with CHB

Previous studies reported that antibodies against PreS1 can inhibit binding of HBV particles to the NTCP receptor and neutralize infection (10). However, only a small percentage (5-7.5%) of individuals with CHB develop anti-PreS1 responses (15, 16). To confirm previous observations, we performed anti-PreS1 endpoint ELISA assays with sera from 61 treatment naïve CHB patients, with each of the five major HBV genotypes (A-E) represented in the cohort (Figure 1A). Consistent with previous reports, only 6 of 61 (9.8%) CHB patients had detectable anti-PreS1 antibodies above background levels (Figure 1B), with genotypes A, B and E represented amongst all the responders, despite using a genotype D serotype ayw PreS1 in the assay. Alignment of the PreS1 amino acid sequences from genotypes A to E shows a high degree of conservation between the genotypes (Figure 1C) with 100% identity in the NTCP receptor binding domain (residues 9 to 18 in bold red (8, 11), genotype D numbering). These data suggest that anti-PreS1 antibodies raised against one genotype may neutralize other HBV genotypes. In summary, our data confirm that anti-PreS1 antibody titers are low in CHB, suggesting a therapeutic opportunity to induce, restore or rescue this response in chronic disease.

**Figure 1.**
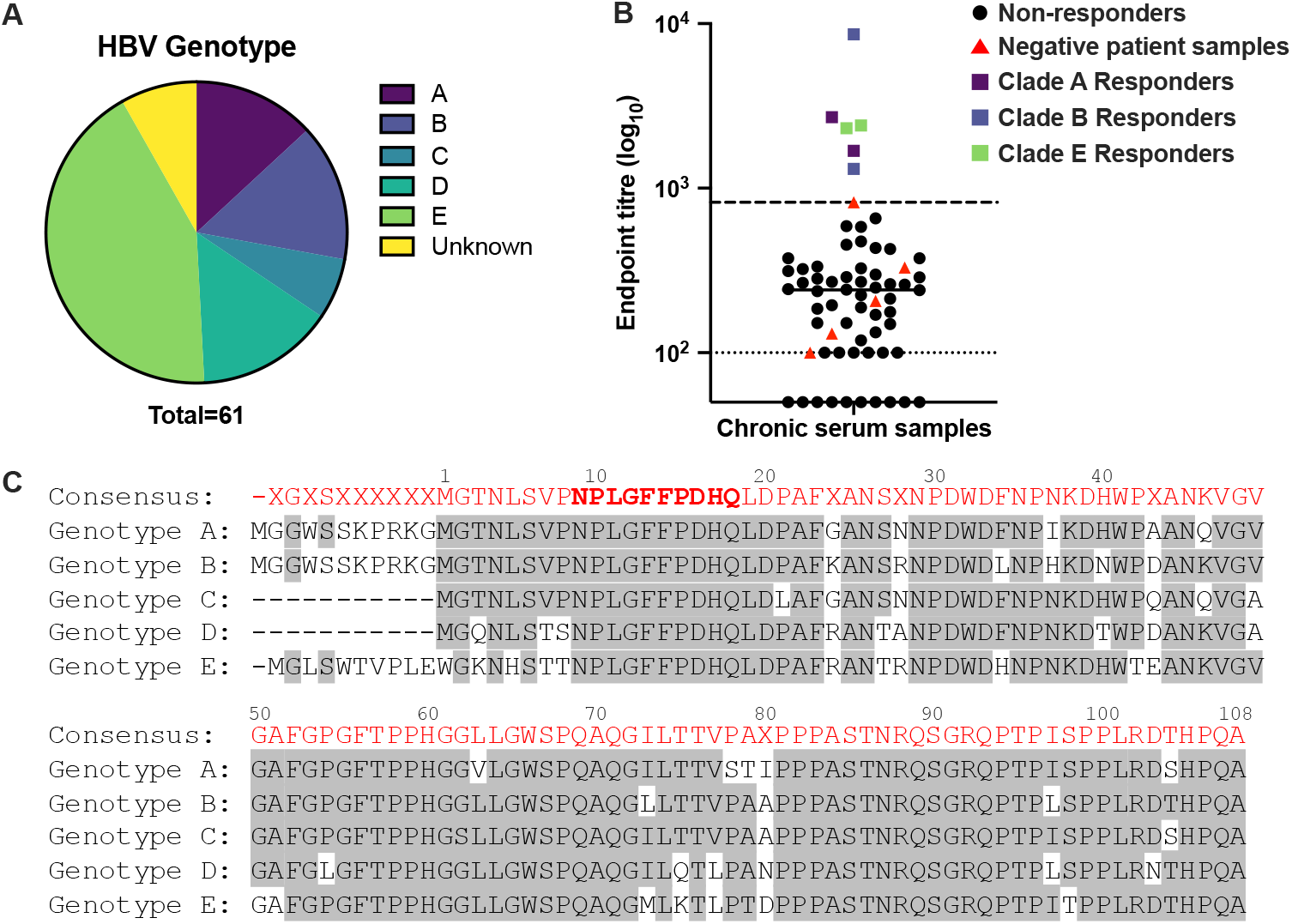
Anti-PreS1 antibody responses are rarely observed in individuals suffering from chronic hepatitis B virus (CHB) **A)** Distribution of genotypes found in CHB samples tested in B. **B)** Levels of anti-PreS1 antibodies were measured by endpoint ELISA in serum from 61 CHB patients and 5 HBV negative patients. Red triangles = Negative patients; Black circles = CHB patients with no anti-PreS1 antibodies (non-responders); Purple, blue and green squares = CHB patients with anti-PreS1 antibodies (responders). The different colours correspond to the different patient genotypes using the same colour scheme as in A. Lower dotted line = limit of detection (LOD), interpolated values below the LOD were reassigned the value of the LOD, uninterpolatable values were assigned an arbitary value of 50 in order to plot them on the graph. Higher dashed line = upper limit of negative patients. **C**) Alignment of the PreS1 amino acid sequence from genotypes A to E. Amino acids highlighted in light grey match the consensus sequence in red. The sequence in bold red from amino acids 9 to18 (counting based on genotype D serotype ayw) is the NTCP-binding domain (Yan et al 2012, Glebe et al 2005).

### Extensive coverage of recombinant Ad particles with PreS1 can be achieved without reducing vector infectivity

We exploited DogTag/DogCatcher protein superglue to decorate recombinant Ad vector particles with PreS1 antigen. DogTag (23 amino acids) was genetically inserted into surface exposed loops on the Ad capsid (hexon hypervariable loop 5)(32) and DogCatcher (15 kDa) fused to the N terminus of PreS1 (genotype D, ayw) for expression as a recombinant protein. Upon co-incubation, hexon-DogTag on adenoviral particles and DogCatcher-PreS1 recombinant protein formed a spontaneous isopeptide bond, resulting in covalent binding of PreS1 to the Ad particles (Figure 2A). Increasing concentrations of DogCatcher-PreS1 resulted in a dose-dependent increase in the coupling efficiency up to a maximum of 70% of virion-associated hexon protein (Figure 2B and C). Additional ligand did not increase coverage beyond 70%, presumably due to steric hinderance preventing the reactivity of the remaining free DogTag loops. Nonetheless, since hexon is the major Ad capsid protein, with 720 copies per virion, a coupling efficiency of 70% corresponds to ∼500 copies of PreS1 per particle. Importantly, even at the highest levels of particle coverage, we observed no reduction in the ability of PreS1-decorated Ad vectors to transduce HEK293 cells *in vitro* compared with an undecorated Ad (Figure 2C).

**Figure 2.**
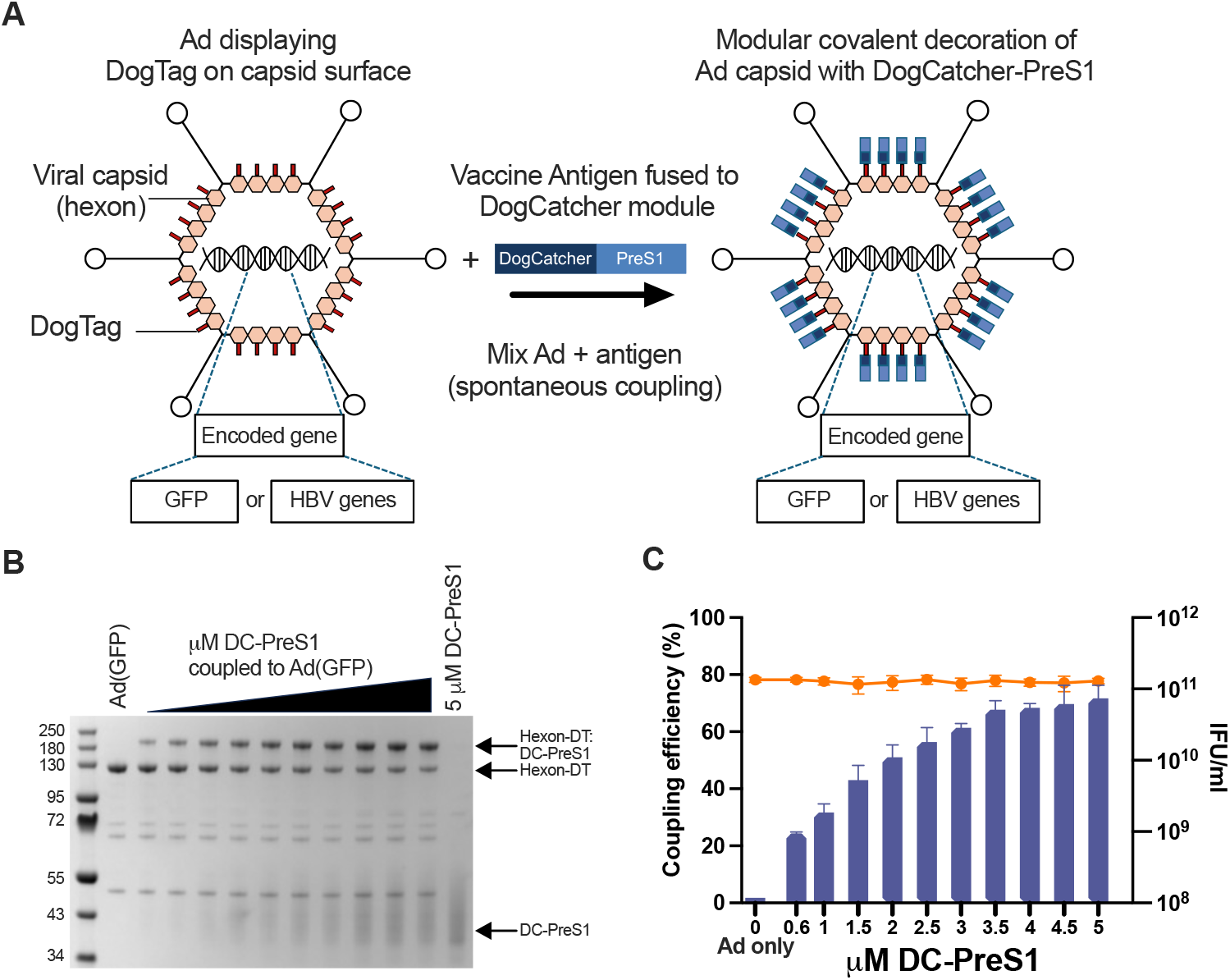
HBV PreS1 can be efficiently coupled to the Ad hexon protein with no loss of Ad infectivity. **A**) Display of DogCatcher-fused PreS1 peptide on the surface of the Ad capsid achieved via covalent attachment to DogTag inserted into hexon surface loop 5 (HVR5). **B**) Representative SDS-PAGE and Coomassie blue staining analysis and **C**) Coupling efficiency (% hexon coverage; blue bars) and infectivity (IFU/ml; orange circles) of Ad-DogTag virions (1E+10 viral particles), incubated with an increasing concentration of DogCatcher-PreS1 protein (0.6 – 5 μM) at 4 °C for 16 h. Samples were analysed in duplicate and the data show mean ± SD of triplicate independent repeats.

### PreS1-decorated Ad elicits anti-PreS1 serum antibody responses

To assess immunogenicity of PreS1-decorated Ad we immunized C57BL/6 mice with GFP encoding Ad vectors, either undecorated or with low (30%, Ad:PreS1^low^) or high (70%, Ad:PreS1^high^) PreS1 capsid coverage. A further group of animals were immunized with DogCatcher-PreS1 recombinant protein alone (at an equivalent PreS1 protein dose to Ad:PreS1^high^ vaccinated animals), formulated in a squalene-based adjuvant (AddaVax) (Figure 3A). Three weeks after a single vaccine dose, immunoglobulin (Ig) G antibody responses against PreS1 were generated in most animals vaccinated with PreS1-decorated Ad (6/6 Ad:PreS1^high^, 4/6 Ad:PreS1^low^) but no detectable responses were achieved after vaccination with DogCatcher-PreS1 protein in adjuvant (Figure 3B). Two weeks after a homologous boost, anti-PreS1 IgG responses further increased in animals vaccinated with PreS1-decorated Ad, with robust responses achieved in all animals. In contrast, only 4/6 animals vaccinated with DogCatcher-PreS1 protein achieved detectable responses post boost, and responses were significantly lower than the Ad:PreS1 vaccinated animals (Figure 3B). As expected, no anti-PreS1 responses were generated in animals vaccinated with the undecorated Ad(GFP) vector. In the same experiment, we assessed the impact of Ad capsid decoration on cellular immune responses against GFP. No differences in the magnitude of splenic CD8+ T-cell responses (as measured by IFNγ-ELISPOT against a C57BL/6 CD8+ T-cell epitope EGFP_118-126_) were observed between PreS1-decorated and undecorated Ad (Figure 3C). These data demonstrate that PreS1 capsid decoration does not impair cellular immune responses against the encoded transgene, consistent with our previous observation that PreS1 decoration does not impair Ad transduction *in vitro*.

**Figure 3.**
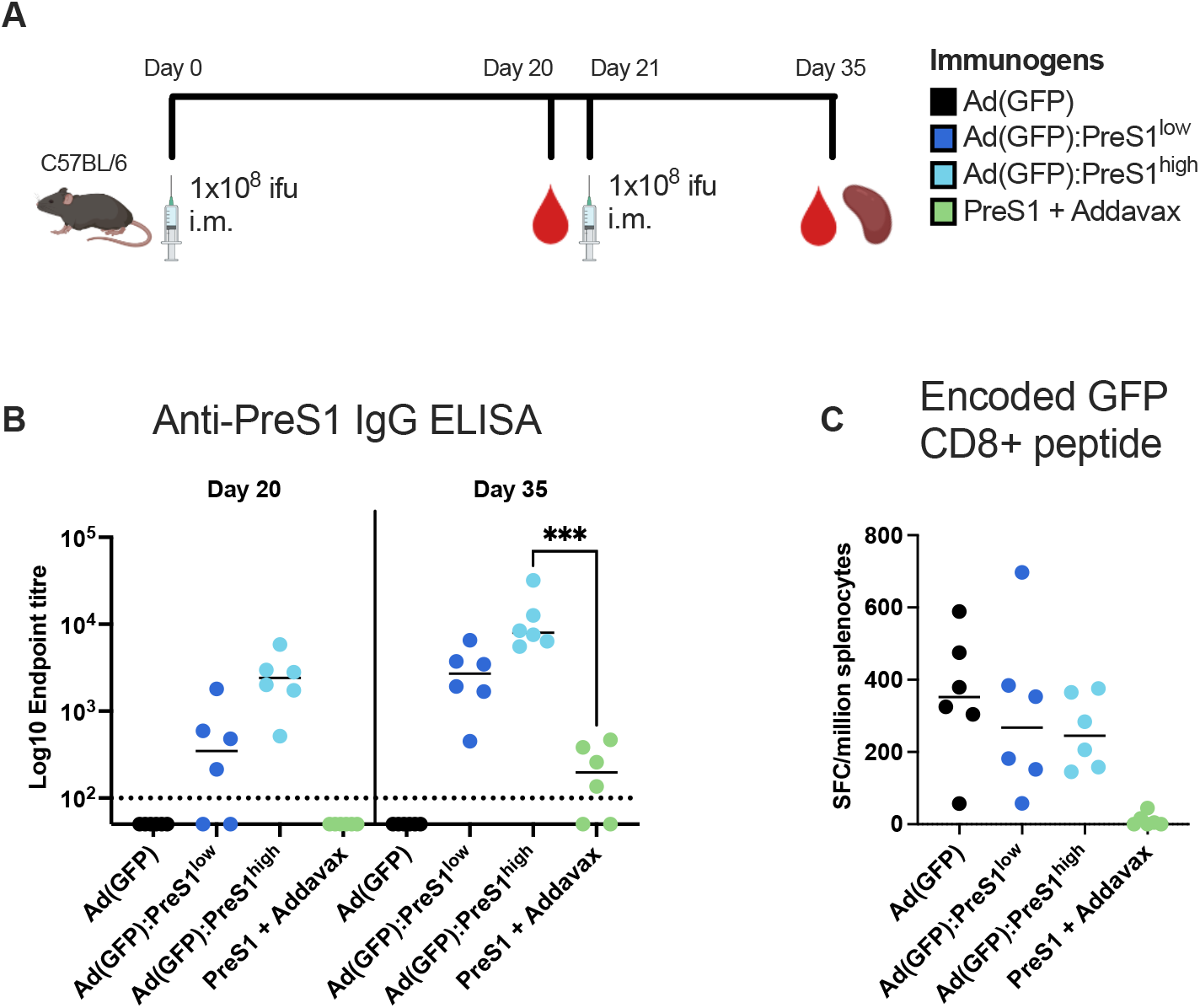
Anti-PreS1 antibodies are generated by capsid display of PreS1 peptide while maintaining encoded antigen CD8+ T-cell responses. **A**) Schematic immunization schedule. C57BL/6 mice (six per group) were immunized intramuscularly (i.m.) in homologous prime boosts regimens with 1E+8 infectious units/mouse of undecorated Ad(GFP), Ad(GFP) with either low (30%, PreS1^low^) or high (70%, PreS1^high^) PreS1 capsid coverage, or DC-PreS1 protein in Addavax (0.2 μg). **B**) Serum IgG antibody responses to PreS1 at D20 and D35 measured by endpoint ELISA. Dotted line represents limit of detection. Uninterpolated samples were given an arbitrary value of 50. Titers at D35 in PreS1-containing vaccine groups were stastically compared. **C**) IFNγ-ELISpot responses (spot forming cells (SFC)/million splenocytes) in spleens at D35 against a GFP CD8+ T-cell epitope peptide, DTLVNRIEL. Ad vaccinated groups were stastically compared. For all graphs, median responses are shown and statistical analyses were determined by Kruskal-Wallis with Dunn’s test for multiple comparisons, ***p < 0.001; non significant comparisons are not shown.

### A PreS1-decorated Ad encoding HBV antigens induces anti-viral humoral and cellular immunity

In parallel, Ad vectors encoding various arrangements of L-HBsAg, S-HBsAg and HBcAg were designed and produced (Figure S1A). Expression of HBV antigens from Ad-transduced HEK293 cells was confirmed by western blotting (Figure S1B). Multiple L-HBsAg and S-HBsAg bands were detected, likely due to different glycosylation patterns (40). To enable expression of multiple antigens to be driven by a single promoter, we utilized the 2A sequence from Foot and Mouth Disease Virus to promote ribosome skipping (41).

An immunogenicity study in C57BL/6 mice was performed to compare Ad vectors encoding HBcAg and S-HBsAg antigens, separated by a 2A sequence (C_2A_S), with Ad encoding L-HBsAg (L). Ad(C_2A_S) and Ad(L) vectors were either undecorated or decorated with PreS1 at ∼30% capsid coverage (comparable to PreS1^low^ in Figure 3). Immunization was performed following the same homologous prime-boost schedule as previously described in Figure 3A. Serum anti-PreS1 IgG responses were measured by ELISA at day 20 and day 35 (Figure 4A). As expected, Ad vectors only elicited detectable PreS1 antibody responses if they encoded L or were decorated with PreS1. By day 35, both PreS1-decorated Ads induced higher PreS1 antibody titers than undecorated Ad(L), with the highest titers induced by PreS1-decorated Ad encoding L, Ad(L):PreS1 (Figure 4A). CD8+ T-cell responses in vaccinated animals were assessed by spleen IFNγ-ELISPOT, with peptide stimulation using previously characterised C57BL/6 CD8+ T-cell epitopes from L/S and C antigens (Figure 4B). Robust CD8+ T-cell responses against L/S were detected in all animals, with Ad vectors encoding L eliciting stronger responses than vectors encoding C_2A_S. PreS1 decoration had no effect on the magnitude of T-cell responses (Figure 4B). Similarly, strong CD8+ T-cell responses against C were elicited in both groups vaccinated with Ad(C_2A_S), with response magnitudes comparable between PreS1-decorated and undecorated vectors (Figure 4B). Antigen-specific T-cell cytokine responses were assessed by intracellular cytokine staining (ICS) using pools of overlapping peptides covering full-length L-HBsAg (L) and HBcAg (C) (Figure 4C). The gating strategy is shown in Figure S2, and phorbol 12-myristate 13-acetate/ionomycin stimulation was used as a positive control (Figure S3). In agreement with the IFNγ-ELISPOT data, PreS1 decoration had no effect on the magnitudes of CD8+ T-cell cytokine responses, and Ad(L) induced higher L-specific IFNγ and TNFα responses than Ad(C_2A_S) (Figure 4C, S3). L- and C-specific IFNγ+CD4+ T-cell responses were also detected, albeit at slightly lower frequencies than the IFNγ+CD8+ responses (Figure 4C), but CD4+ IL-2 and TNFα responses were minimal (Figure S3).

**Figure 4.**
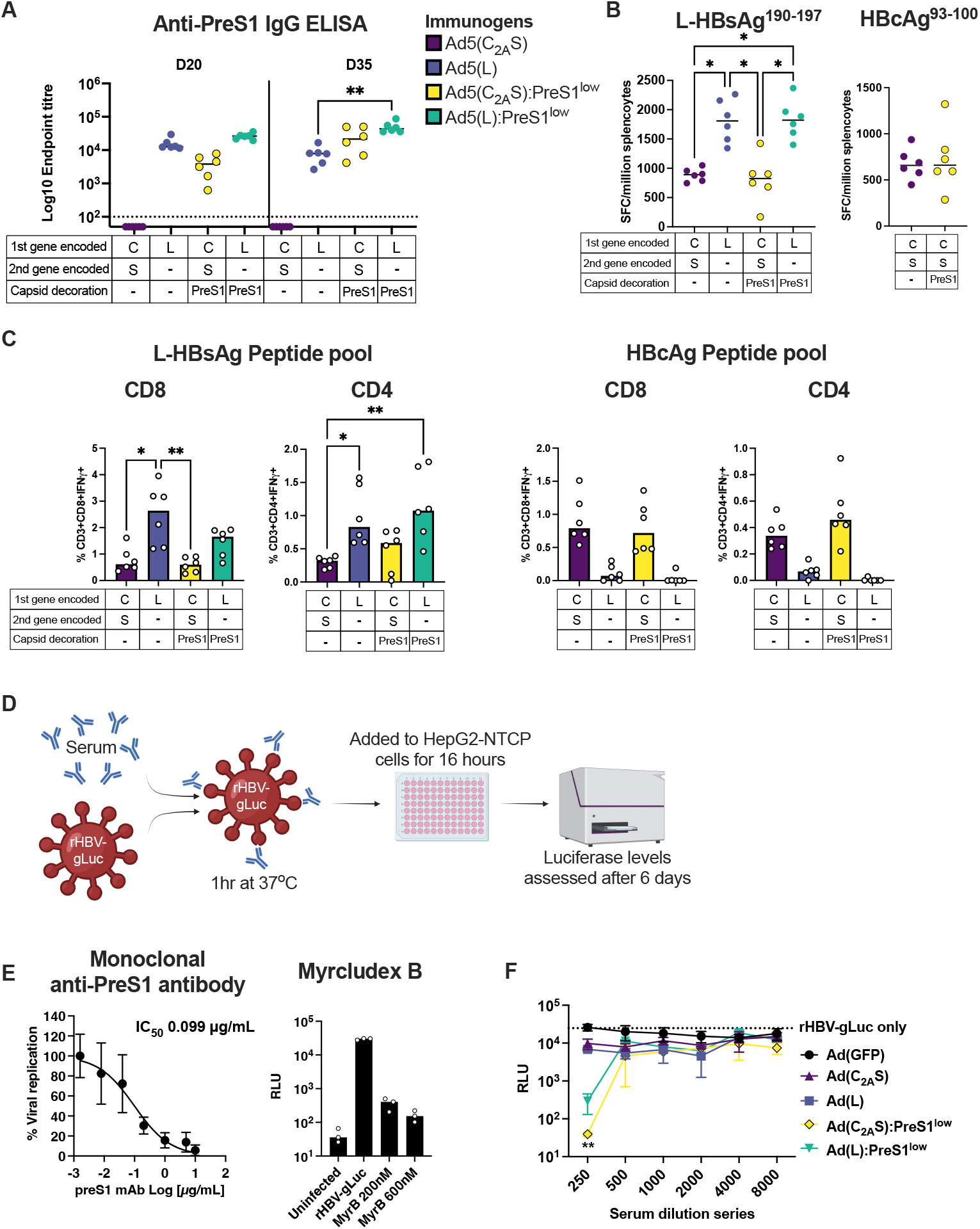
Optimal anti-PreS1 antibody responses and anti-L T-cell responses produced by vectors decorated with PreS1 and encoding L. C57BL/6 mice (six per group) were immunized intramuscularly (i.m) with Ad vectors encoding HBcAg and S-HBsAg or L-HBsAg with or without PreS1 peptide decoration in homologous prime boost regimens (1E+8 infectious units/mouse). Vaccine immunogens in each group are detailed in panel A and below each graph. **A**) Serum IgG antibody responses to PreS1 at D20 and D35 measured by endpoint ELISA. Dotted line represents limit of detection. Uninterpolated samples were given an arbitrary value of 50. Titers at D35 in PreS1-containing vaccine groups (encoded or surface displayed) were stastically compared. **B**) IFNγ-ELISpot responses (spot forming cells (SFC)/million splenocytes) in spleens at D35 against an L-HBsAg CD8+ T-cell epitope peptide, VWLSVIWM or a HBcAg CD8+ T-cell epitope peptide, MGLKFRQL. **C**) Percentages of IFNγ-producing CD3+ CD8+ T-cells and CD3+ CD4+ T-cells at D35 responding to L-HBsAg and HBcAg peptide pools. For the HBcAg peptide pools only groups encoding HBcAg were statistically compared. **D)** Schematic describing the HBV neutralization assay. **E**) Neutralization of rHBV-gLuc infection using either a dilution series of a mouse monoclonal anti-PreS1 antibody or two concentrations of MyrcludexB (Bulevirtide™), RLU = Relative Light Units. **F**) Neutralization of rHBV-gLuc with heat-inactivated pooled serum from mice immunized with Ad vectors encoding HBV genes with and without PreS1 capsid decoration. Experiments were performed in technical triplicates with 3 independent repeats. At the 1:250 dilution, groups were compared to Ad(GFP) control. For graphs A-C and F median responses are shown, for E and G mean values with SD are shown. Stastical analyses were carried out on data in A-C, and F by Kruskal-Wallis with Dunn’s test for multiple comparisons where more than one comparison was made and by Mann-Whitney for a single comparison, *p < 0.05; **p < 0.01; ***p < 0.001; non-significant comparisons are not shown.

As the PreS1 region of L-HBsAg is essential for HBV entry into cells, we hypothesised that anti-PreS1 antibodies induced by these vaccines would neutralize virus infection. We established an *in vitro* HBV neutralization assay using Gaussia Luciferase-expressing reporter HBV (rHBV-gLuc) for *de novo* infection of hepatoma cells (Figure 4D). This virus encodes gaussia luciferase in place of the viral polymerase but uses authentic HBV entry mechanisms with PreS1 engagement with NTCP to infect cells and establish covalently closed circular DNA in a single cycle infection. Incubation of rHBV-gLuc with either an anti-PreS1 monoclonal antibody or the peptide entry inhibitor, MyrcludexB (Bulevirtide™, Hepcludex™), successfully inhibited infection of HepG2-NTCP cells (Figure 4E). MyrcludexB is a myristolated lipopeptide comprising 47aa (2-48) of the PreS1 region which binds NTCP, thereby blocking HBV and HDV cell entry. Serum (at day 35) from mice immunized with undecorated Ad(C_2A_S) or Ad(L) showed only weak HBV neutralizing activity (2.7 and 3.8 fold inhibition respectively, relative to control Ad(GFP) at 1:250 serum dilution), while PreS1-decorated Ad(C_2A_S):PreS1 and Ad(L):PreS1 elicited serum responses that neutralized HBV at the lowest dilution (662-fold and 89-fold inhibition respectively at 1:250 dilution) (Figure 4F). These data demonstrate that PreS1 capsid decoration is required to induce serum neutralizing antibodies.

### Optimizing anti-HBV immunity with PreS1-decorated Ad

To further improve immunogenicity, Ad vectors encoding L, L_2A_C and C_2A_L (L and C antigens in the opposite order) were decorated with PreS1 at 70% capsid coverage (comparable to PreS1^high^ in Figure 3) rather than 30% as in the previous experiment and C57BL/6 mice were immunized following the same dosing regimen. Robust anti-PreS1 IgG ELISA titers were achieved in all three groups and were comparable between groups (Figure 5A). CD8+ T-cell responses against an L/S epitope, as assessed by IFNγ-ELISPOT, were comparable between all three groups demonstrating that the addition of C and the order of L and C antigens had a minimal effect on the magnitude of the T-cell responses (Figure 5B). Similar responses were noted against the C epitope between Ad(L_2A_C):PreS1 and Ad(C_2A_L):PreS1 vaccinated groups (Figure 5B). CD8+ and CD4+ T-cell cytokine responses against L were comparable between all three groups (Figure 5C and Figure S4). CD8+ and CD4+ T-cell cytokine responses against the C peptide pool were comparable between Ad(L_2A_C):PreS1 and Ad(C_2A_L):PreS1 vaccinated groups, though there was a non-significant trend towards higher IL-2+CD8+ T-cells in Ad(L_2A_C):PreS1 vaccinated animals (Figure S4).

**Figure 5.**
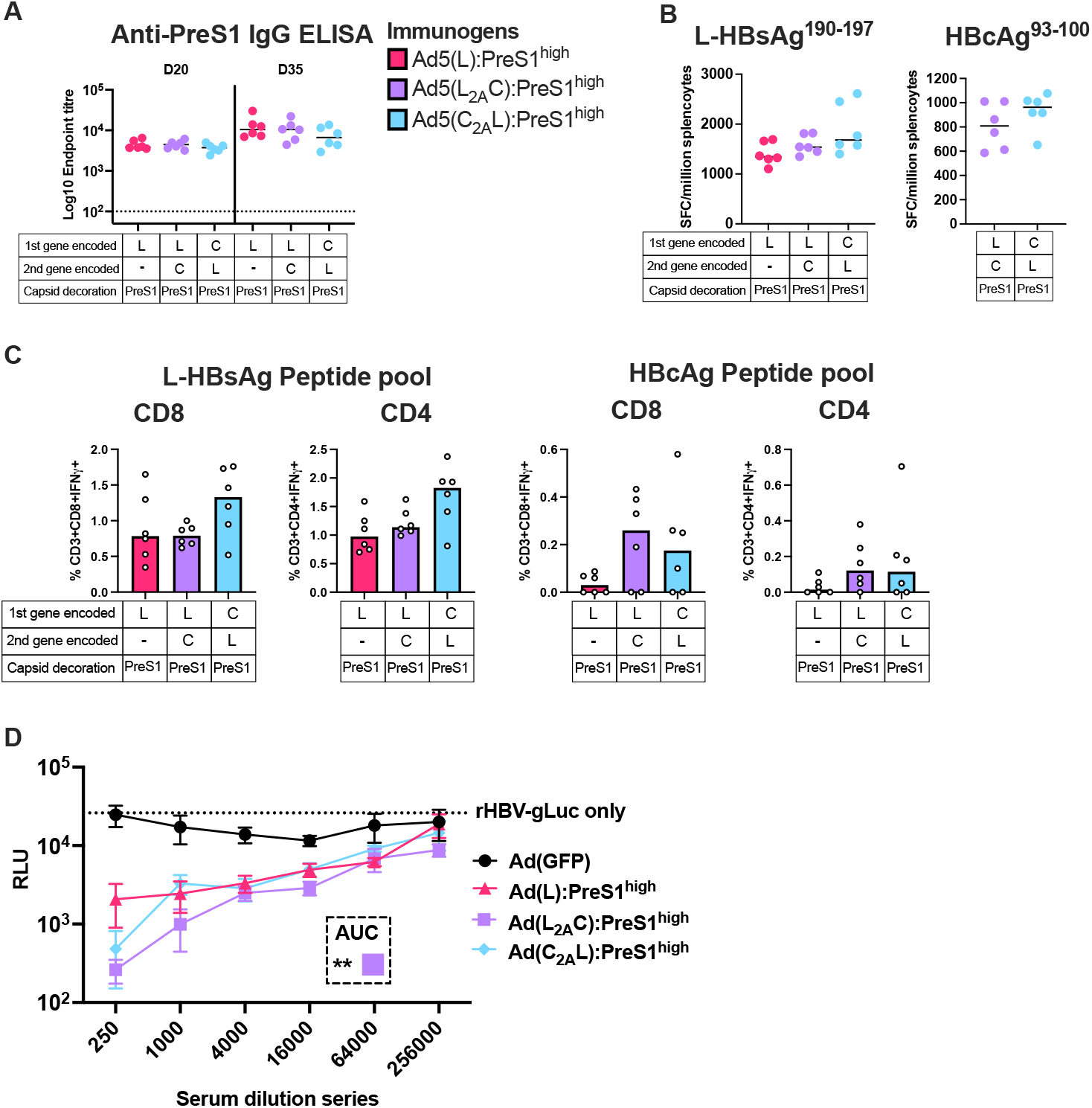
Displaying high levels of PreS1 on the Ad capsid surface enhances HBV neutralization. C57BL/6 mice (six per group) were immunized intramuscularly (i.m) with PreS1^high^ decorated Ad encoding L-HBsAg or L-HBsAg and HBcAg, in homologous prime boost regimens (1E+8 infectious units /mouse). Immunogens used are detailed in panel A and below each graph. **A**) Serum IgG antibody responses to PreS1 at D20 and D35 measured by endpoint ELISA. Dotted line represents limit of detection. D35 groups were stastically compared. **B**) IFNγ-ELISpot responses (spot forming cells (SFC)/million splenocytes) in spleens at D35 against an L-HBsAg CD8+ T-cell epitope peptide, VWLSVIWM or a HBcAg CD8+ T-cell epitope peptide, MGLKFRQL. **C**) Percentages of IFNγ-producing CD3+ CD8+ T-cells and CD3+ CD4+ T-cells at D35 responding to L-HBsAg and HBcAg peptide pools. For HBcAg peptide pools only groups immunized with vaccines encoding HBcAg were stastically compared. **D**) Neutralization of rHBV-gLuc with heat-inactivated pooled serum from immunized mice. Experiments were performed in technical triplicates with 3 independent repeats. Area under the curve (AUC) was determined for all groups and stastically compared to Ad(GFP). For graphs A-C median responses are shown, for D mean values with SD are shown. Stastical analyses were carried out on data in A-C and AUC of D by Kruskal-Wallis with Dunn’s test for multiple comparisons where more than one comparison was made and by Mann-Whitney for a single comparison, non-significant comparisons are not shown.

Serum HBV neutralization was assessed as described previously. Mice vaccinated with Ad(L):PreS1, Ad(L_2A_C):PreS1 and Ad(C_2A_L):PreS1 showed neutralizing activity (Figure 5D), and in contrast to the previous experiment, neutralization was maintained upon serial dilution of the serum up to a dilution of at least 1:16,000 suggesting a more robust response. ID_50_ values were 120956 for Ad(L):PreS1, 264763 for Ad(L_2A_C):PreS1, and 119051 for Ad(C_2A_L):PreS1. The strongest serum neutralization was observed in the Ad(L_2A_C):PreS1 vaccinated group, that showed statistically significant neutralization compared to the control group by area under the curve analysis (Figure 5D). Our data suggest that PreS1 decoration at a higher capsid coverage (70% vs 30% in the previous experiment) induced more potent neutralizing responses.

### PreS1-decorated Ad achieves serum neutralization of Hepatitis D virus

HDV is a small defective RNA virus that can use the HBV encoded surface glycoproteins to assemble infectious particles (42) and frequently co-infects with HBV. MyrcludexB has been shown to induce sustained virologic and biochemical responses in chronic HDV (43) and is approved in the EU for the treatment of HDV-associated compensated liver disease. Given that both HDV and HBV are reliant on PreS1 engagement with NTCP to infect cells, we hypothesised that vaccine-induced anti-PreS1 antibodies should neutralize HDV infection of HepG2-NTCP cells. To test this, we incubated HDV with pooled serum from mice immunized with Ad(HBV L_2A_C):PreS1 (from the experiment shown in Figure 5). In parallel, we incubated HDV with an anti-PreS1 monoclonal antibody or MyrcludexB as controls, as per our HBV neutralisation assay. Complexes were added to HepG2-NTCP cells, and HDV infection assessed by immunostaining for HDV antigen at 6 days post infection (Figure 6A). We observed a marked reduction in HDV infected cells after exposure to serum from mice immunized with Ad(HBV L_2A_C):PreS1 (Figure 6B). This phenocopied the results observed with the control anti-PreS1 (10 μg/ml) and MyrcludexB (200 nM). These data show that PreS1-decorated Ad may have additional utility in chronic HBV-HDV co-infection, due to the homologous cellular entry mechanisms used by these two pathogens.

**Figure 6.**
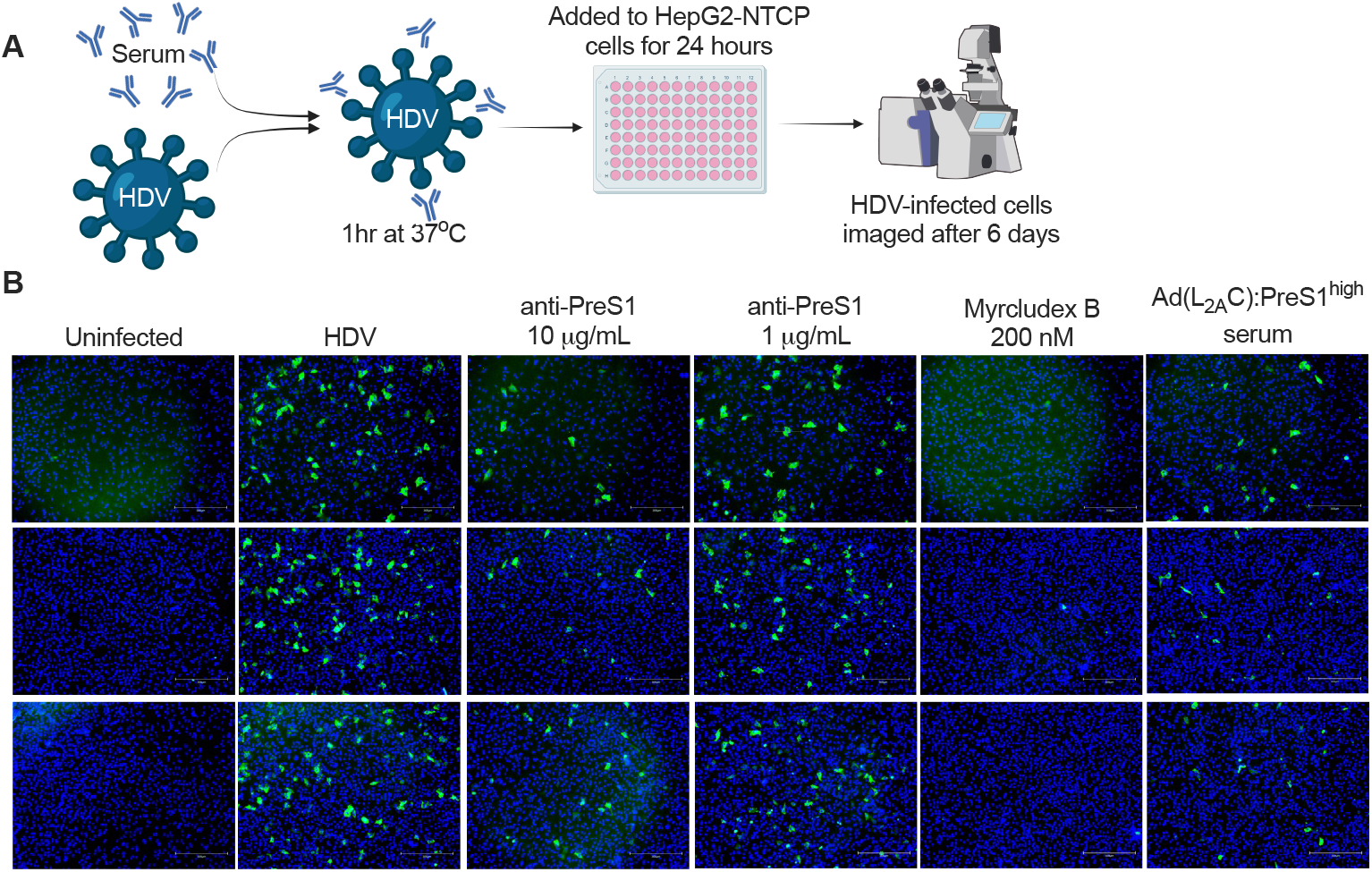
Serum from mice immunized with PreS1-decorated Ad neutralizes HDV. **A**) Schematic of the HDV neutralization assay. **B**) Images of HepG2-NTCP cells either uninfected or infected with HDV in the presence or absence of anti-PreS1 antibodies, Myrcludex B or serum from mice immunized with Ad(L_2A_C):PreS1^high^ (from the experiment shown in Figure 5). Images from 3 independent infected wells are shown. Scale bar - 300 μm

## Discussion

In this study, we describe a novel therapeutic vaccination strategy to induce neutralizing antibody responses and potent T-cell immunogenicity against HBV in a single platform using a PreS1-decorated Ad vector encoding multiple HBV antigens. PreS1 is required for HBV and HDV entry into cells (10), and hence PreS1-targeted neutralizing antibodies could prevent *de novo* infection, limiting the spread of infection and associated intrahepatic inflammation (44). We confirmed that anti-PreS1 immunity is limited or absent in CHB (Figure 1B), presenting an opportunity for therapeutic intervention.

Display of peptide and protein antigens on nanoparticles is an effective strategy to improve antigen-specific antibody titers upon vaccination (45). Here, we utilized DogTag/DogCatcher protein superglue to decorate recombinant Ad particles with PreS1 (Figure 2). PreS1-decorated Ad achieved significantly higher anti-PreS1 IgG titers compared to recombinant DogCatcher-PreS1 protein in adjuvant (Figure 3B) or a conventional undecorated Ad encoding L-HBsAg (Figure 4A). Crucially, PreS1-decoration was essential for the induction of neutralizing activity (Figure 4F) and highly decorated particles induced a more potent response (Figure 5D, ID_50_ >10^5^). A key advantage of PreS1-decorated Ad over protein nanoparticle and virus-like-particle technologies for therapeutic vaccination is that Ad vector vaccines induce potent T-cell immunity, particularly CD8+ T-cell responses, against antigens encoded in the vector genome. PreS1-decorated Ad induced robust CD8+ and CD4+ T-cell responses against encoded HBV antigens, L-HBsAg and HBcAg (Figure 4,5). Bivalent cassettes induced comparable T-cell responses to a monovalent cassette expressing only L-HBsAg, indicating that co-expression of HBcAg did not lower responses against L-HBsAg through antigenic competition. Resolution of acute infection typically requires CD8+ T-cell responses against multiple antigens (22) so multivalent antigen cassettes are likely to be advantageous. Despite achieving promising immunogenicity in HBV naïve mice, it remains to be established whether our strategy can overcome immune tolerance in CHB. Therapeutic strategies using recombinant viral vectors (including Ad) and PreS1-decorated nanoparticles, have demonstrated encouraging immunogenicity in immune tolerant HBV-carrier mouse models (13, 17, 28) and early-stage clinical studies in CHB patients (25).

Establishing a functional cure for CHB will almost certainly require a combinatorial approach using multiple therapeutic components targeting different stages of the viral life cycle. Feld and colleagues highlighted three principal strategies; i) inhibition of viral replication to prevent further spread of infection (e.g. entry inhibitors or capsid inhibitors) ii) antigen reduction to alleviate immune tolerance (e.g. siRNAs) and iii) immune modulation to re-establish anti-HBV immunity (e.g. checkpoint inhibitors and therapeutic vaccines) (46). PreS1-decorated Ad addresses two of these key aspects and could contribute towards the development of a functional cure in combination with translation inhibitors, checkpoint inhibitors, and / or complimentary vaccination strategies (e.g., heterologous prime-boost regimens). Several combination approaches are currently being assessed in clinical trials. A PreS-containing HBsAg virus-like-particle vaccine (BRII-179) (comprising L-, M- and S-HBsAg isoforms) co-administered with an siRNA (Elebsiran) to lower circulating antigen levels and reduce immune tolerance, effectively re-established anti-HBV immunity including anti-HBs antibodies and PreS1/PreS2-specific CD4+ T-cell responses (47). Such an approach could suppress HBsAg and HBV DNA once siRNA treatment is withdrawn. Indeed, one of the participants in the vaccinated arm of this study met the criteria for NRTI discontinuation. The authors hypothesized that anti-HBV neutralizing antibodies and PreS1- and PreS2-specific CD4+ T-cells may have helped restrict *de novo* infection and re-establish virological control in this patient (47). BRII-179, a recombinant protein vaccine, did not induce HBV-specific CD8+ T-cell responses. Induction of CD8+ T-cell immunity in this setting, perhaps through the addition of viral vectors, could further improve the virological response to vaccination. In another study, a recombinant Ad vector encoding multiple HBV antigens, ChAdOx1-HBV, was combined with a poxvirus vector, MVA-HBV, in a heterologous prime-boost regimen (VTP-300) to further improve T-cell responses in CHB patients (26). Combination of these therapeutic vaccines with a checkpoint inhibitor (low-dose nivolumab) administered at the time of the MVA boost led to enhanced reduction in HBsAg levels (26). Combining VTP-300 with an siRNA (Imdusiran) also led to improved HBsAg reduction compared to siRNA treatment alone in virally suppressed CHB patients (48).

By targeting PreS1, our approach could provide additional benefit in the context of HBV/HDV co-infection. As expected, serum from mice vaccinated with PreS1-decorated Ad inhibited HDV infection (Figure 5). HDV/HBV co-infection occurs in approximately 5% of CHB patients and is considered the most severe form of viral hepatitis, with more rapid progression to severe liver cirrhosis and hepatocellular carcinoma. Induction of neutralizing antibodies through PreS1 vaccination could provide an attractive alternative or complimentary approach to treatment with currently licenced PreS1-based peptide inhibitors such as Bulevirtide™. Whilst there has been no direct head-to-head comparison, anti-HBs monoclonal antibodies have arguably achieved more substantial declines in HDV RNA compared to Bulevirtide monotherapy (49), perhaps through the additional contribution of Fc-receptor mediated effector functions (50). In a separate study, Bulevirtide combined with pegylated interferon-alpha was more effective at reducing HDV DNA to undetectable levels than Bulevirtide monotherapy (51), further supporting the combination of entry inhibition with immune modulation. Immunological memory acquired through PreS1 vaccination could ultimately provide finite duration treatments for HDV; Bulevirtide currently requires daily subcutaneous administration and relapses frequently occur upon withdrawal (52).

The ability of PreS1-decorated Ad to combine induction of potent HBV/HDV neutralizing humoral immunity with strong cellular immune responses in a single platform differentiates our technology from current state-of-the-art approaches. Conventional viral vector vaccines, including recombinant Ads, do not typically induce strong vaccine-antigen specific B-cell responses, and therefore many viral vector-based therapeutic vaccine studies have neglected to target (or even assess) induction of humoral immunity. Conversely, protein-in-adjuvant vaccines may have lacked activity in CHB due to weak or absent CD8+ T-cell induction. Further development of our platform could include capsid display of additional neutralizing epitopes (e.g. from PreS2 or the S-HBsAg antigenic loop) and / or incorporation of more encoded antigens (e.g. polymerase) to increase the breadth of T-cell immunity (25). Combining PreS1-targeted entry inhibition with robust anti-HBV/HDV T-cell induction offers a promising strategy to improve therapeutic outcomes in chronic HBV and HDV infection.

## Abbreviations

Ad: Adenovirus
CHB: Chronic hepatitis B virus
HBcAg: Hepatitis B virus core antigen
Immunoglobulin: Ig
L-HBsAg: Hepatitis B virus large surface antigen
MVA: Modified vaccinia Ankara
NA: Nucleos(t)ide analogues
NTCP: Sodium taurocholate co-transporting polypeptide
S-HBsAg: Hepatitis B virus small surface antigen

## Acknowledgements

The authors would like to thank Rosa D’Agostino, Laura Bilbé, Lesley Bowman, Jeanette Wagener, David Bitto, Sophie Porret, Antonia Hook, Laura Silva-Reyes and Jakub Kopycinski for helpful advice and assistance. We thank Viv Clark, Heather Chandler, Stephen Laird, Douglas dos Santos Passos, and Luke Harris for animal husbandry and technical assistance. We also thank Stephan Urban (University of Heidelberg) for supplying HepG2-NTCP cells and Jochen Wettengel and Ulrike Protzer (Technical University of Munich) for purified HB-Luciferase reporter virus.

## Notes

**Financial support and sponsorship** Research in the McKeating laboratory is funded by the Wellcome Trust Discovery Award 225198/Z/22/Z) and the Chinese Academy of Medical Sciences (CAMS) Innovation Fund for Medical Science (China 2018-I2M-2-002). PACW was supported by the Chinese Academy of Medical Sciences (CAMS) Innovation Fund for Medical Science (China 2018-I2M-2-002)

### Competing Interest Statement

R.A.R, L.M.R, B.G.C, and M.D.J.D are employees of SpyBiotech Ltd. S.B. is CSO and co-founder of SpyBiotech Ltd.

