## Supplemental Material for "PreS1-decorated recombinant adenovirus encoding HBV antigens generates neutralizing humoral and cellular immunity"

**Supplementary methods:**

**Recombinant adenovirus HBV transgene expression**

Transgene expression from HBV-encoding vectors was determined by infecting 293A cells, seeded at 1.25E+6 in 6-well plates, with the Ad vectors at a multiplicity of infection of 100. The next day cells were harvested and 5E+5 cells were lysed in reducing 4x Laemmli sample buffer (Bio-Rad, UK) with 300mM 1,4-dithiothreitol (Merck, UK) at 95°C for 10 minutes. Samples were separated on a 4-12% Bis-Tris gel (Life Technologies, UK) and transferred to a 0.2 μm nitrocellulose membrane (Bio-Rad, UK). Western blots were carried out using the iBind system (Life Technologies, UK). The following antibodies were used: anti-β-Actin rabbit monoclonal antibody (mAb) clone D6A8 at 1:500 (Cell Signaling Technologies, The Netherlands; anti-Ad5 hexon mouse mAb clone 65H6 at 1:500 (Life Technologies, UK); anti-S-HBsAg mouse mAb clone 1837 at 1:200 (Native Antigen, UK); anti-PreS1 mouse mAb clone AP1 at 1:200 (Santa Cruz Biotechnology, USA); anti-PreS2 mouse mAb clone S26 at 1:200 (Santa Cruz Biotechnology, USA); anti-Core mouse mAb clone 10E11 at 1:200 (Santa Cruz Biotechnology, USA). Goat anti-mouse IgG conjugated to alkaline phosphatase (STAR117A at 1:1000, Bio-Rad, UK) was used as a secondary for all mouse primary antibodies and goat anti-rabbit conjugated to alkaline phosphatase (A3687 at 1:1000, Merck, UK) was used as a secondary for the anti-β-Actin rabbit primary antibody. The signal was developed using BCIP/NBT solution (Merck, UK).

**Flow cytometry protocol and antibody cocktails:**

Cells were stained with Fixable Live/Dead Aqua (1:1000 Life Technologies, UK) and Fc receptors were blocked with 2.5 μg/mL CD16/CD32 Fc block (BD Biosciences, UK) before being stained with an extracellular antibody cocktail diluted in FACS wash buffer (PBS with 2 mM EDTA and 0.5% bovine serum albumin) (anti-mouse CD3 APC-Fire750, clone 17A2, 1:40, BioLegend, USA; anti-mouse CD4 PE-Dazzle594, clone GK1.5, 1:200, BioLegend, USA; anti-mouse CD8a BB700, clone 53-6.7, 1:80, BD Biosciences, UK; anti-mouse CD14 BV510, clone Sa14-2, 1:80, BioLegend, USA and anti-mouse CD19 BV510, clone 1D3, 1:200, BioLegend, USA). Cells were fixed in Cytofix/Cytoperm (BD Biosciences, UK) before being stained with an intracellular antibody cocktail diluted in Perm/Wash buffer (BD Biosciences, UK) (anti-mouse IFNγ APC, clone XMG1.2, 1:160, BioLegend, USA; anti-mouse IL-2 PE, clone JES6-5H4, 1:80, BioLegend, USA and anti-mouse TNFα FITC, clone MP6-XT22, 1:400, BioLegend, USA).

**Preparation of HepG2-NTCP cells for the HBV neutralization assay**

For the HBV neutralization assay HepG2-NTCP cells were seeded at 1E+4/well in DMEM (Life Technologies, UK) with 10% Foetal Calf Serum (Gibco, UK), 1% Non-Essential Amino Acids (Life Technologies, UK) and 1% Penicillin & Streptomycin (Life Technologies, UK) (complete medium) in collagen-coated (Sigma) flat-bottomed 96-well plates (Corning, USA). The next day the medium was replaced with complete medium supplemented with 2.5% DMSO (Merck, UK) and the cells incubated for 72 hours. The medium from the HepG2-NTCP cells was removed and replaced with 100 μL complete medium supplemented with 2.5% DMSO and 4% Polyethylene glycol (PEG8000) (Merck, UK). 10 μL of the reporter virus/serum dilution series was added to the cells and the infection allowed to proceed for 16 hours. The viral inocula was removed, cells washed twice with 100 μL PBS, and the medium replaced with 100μL complete medium supplemented with 2.5% DMSO. After 3 days the medium was replenished. After a further 3 days the Gaussia luciferase signal was detected to determine the level of infection.

**Preparation of HDV**

Twenty-four hours prior to transfection, Huh-7 cells were seeded in complete medium at a density of 6E+6 cells/T75 flask. The next day, cells were transfected with 5 μg of pSVLD3 and 5 μg of pT7HB2.7 plasmids using 30 μL of FuGENE-HD reagent (Promega, UK). Following overnight incubation, cells were washed twice with PBS and the medium was replaced with 12.5 mL of complete medium. Supernatant was collected at 3-, 6-, 9-, and 12-days post-transfection, then concentrated to $1/50^{th}$ of the original supernatant volume using Ultra Centrifugal Filter 50 kDa (Merck, UK) and used as HDV inoculum. To quantify genome copy number, RNA from 3 μL of concentrated HDV inoculum was extracted using a Viral RNA Mini Kit (QIAGEN, Germany), and reverse transcribed using a cDNA synthesis kit (PCR Biosystems, UK), according to the manufacturer's protocol (25°C, 10 min; 42°C, 15 min; 48°C, 15 min; 85°C, 10 min). Gene expression was quantified using a SyGreen Blue Mix (PCR Biosystems, UK) using a qPCR program of 95°C, 2 min; 45 cycles of 95°C, 5 sec; 60°C, 30 sec, using primers specific for HDV <5'-TCTTCCTCGGTCAACCTCTT-3' and 5'-ACAAGGAGAGGCAGGATCAC-3'>. Genome copy number was enumerated from a standard curve of pSVLD3 plasmid of known concentration.

**Preparation of HepG2-NTCP cells for the HDV neutralisation assay**

HepG2-NTCP cells were seeded at a density of 1.25E+4 cells/well on collagen coated 96 well plates and cultured in complete medium. Next day the medium was replaced with complete medium containing 2.5% DMSO for a further 72h. Test sera were pre-inoculated with HDV (MOI of 50 genome copies per cell) in the presence of 4% PEG8000 and 2.5% DMSO at 37°C for 1h, then added to the cells for 24h infection. Viral inoculum was removed, and cells washed three times with PBS. Infected cells were maintained with complete medium containing 2.5% DMSO, changing media every 3 days. Six days post infection, cells were fixed with 4% PFA and the number of HDV-infected cells determined by immunofluorescence. Cells were maintained in 5% $CO_2$ and 18% $O_2$.

### Supplemental Figure 1

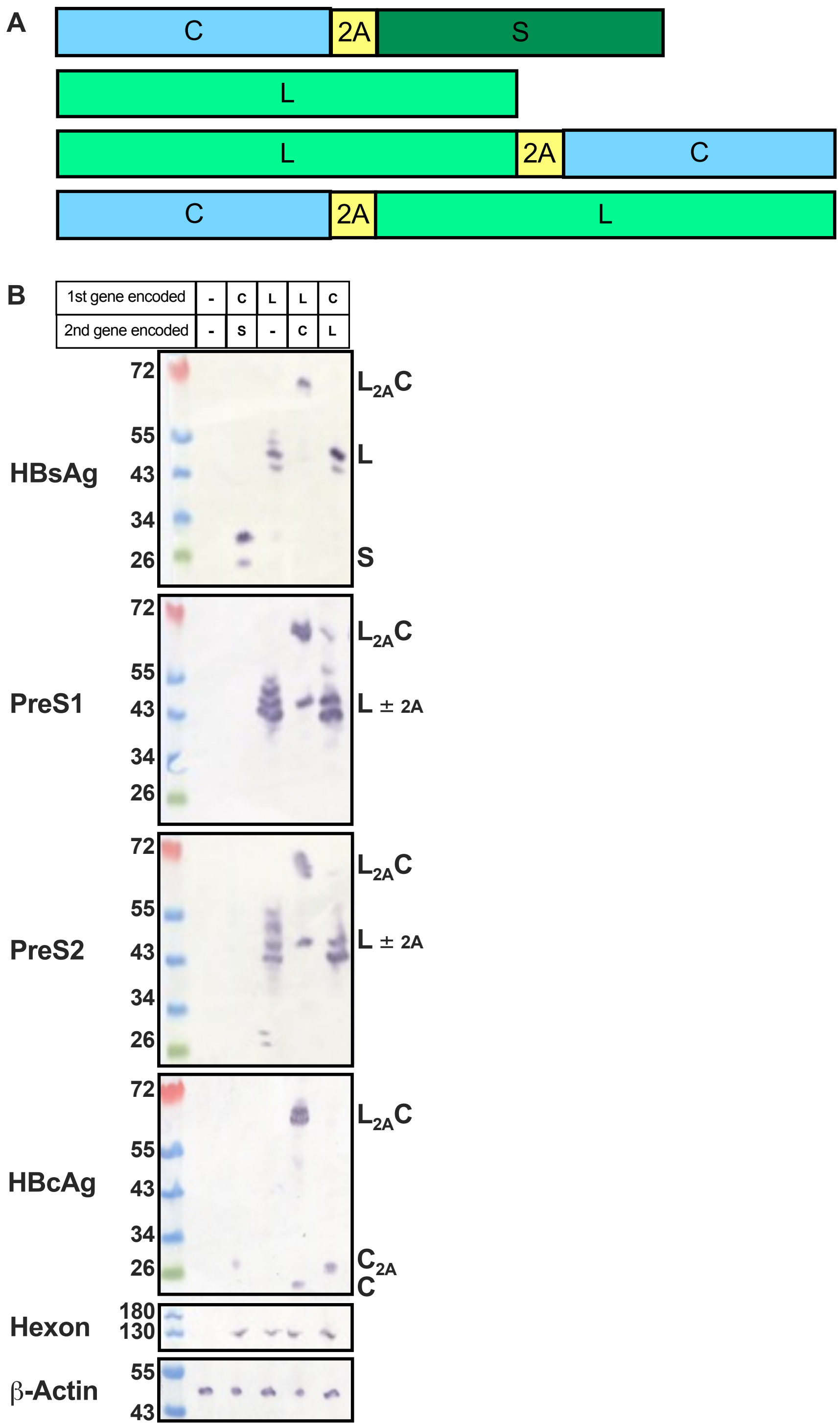


**Supplemental Figure 1 - Encoded HBV genes.**
**A**) Schematic of the different HBV genes encoded in the Ad vectors. Vectors were designed to express HBcAg (C), the S-HBsAg (S) or the L-HBsAg (L), separated, where necessary, by a GSG linker and an F2A sequence. For optimal expression and immunogenicity a Kozak sequence followed by a truncated shark Invariant chain (SIi) and a GSGGSG linker were inserted up stream of the first open reading frame. B) HEK293A cells were infected with Ad vectors at a multiplicity of infection of 100. 24 h post-transfection, cells were lysed and the lysates were analysed by Western blot using mouse anti-HBsAg, mouse anti-PreS1, mouse anti-PreS2 and mouse anti-HBcAg antibodies. Blots probed with mouse anti-Ad5 hexon and rabbit anti-β-actin served as loading controls. Lane 1: cell lysate from un-transduced cells, lanes 2 to 5: cell lysates from cells transduced with Ad encoding HBV genes as detailed in the grid above the first blot.

### Supplemental Figure 2

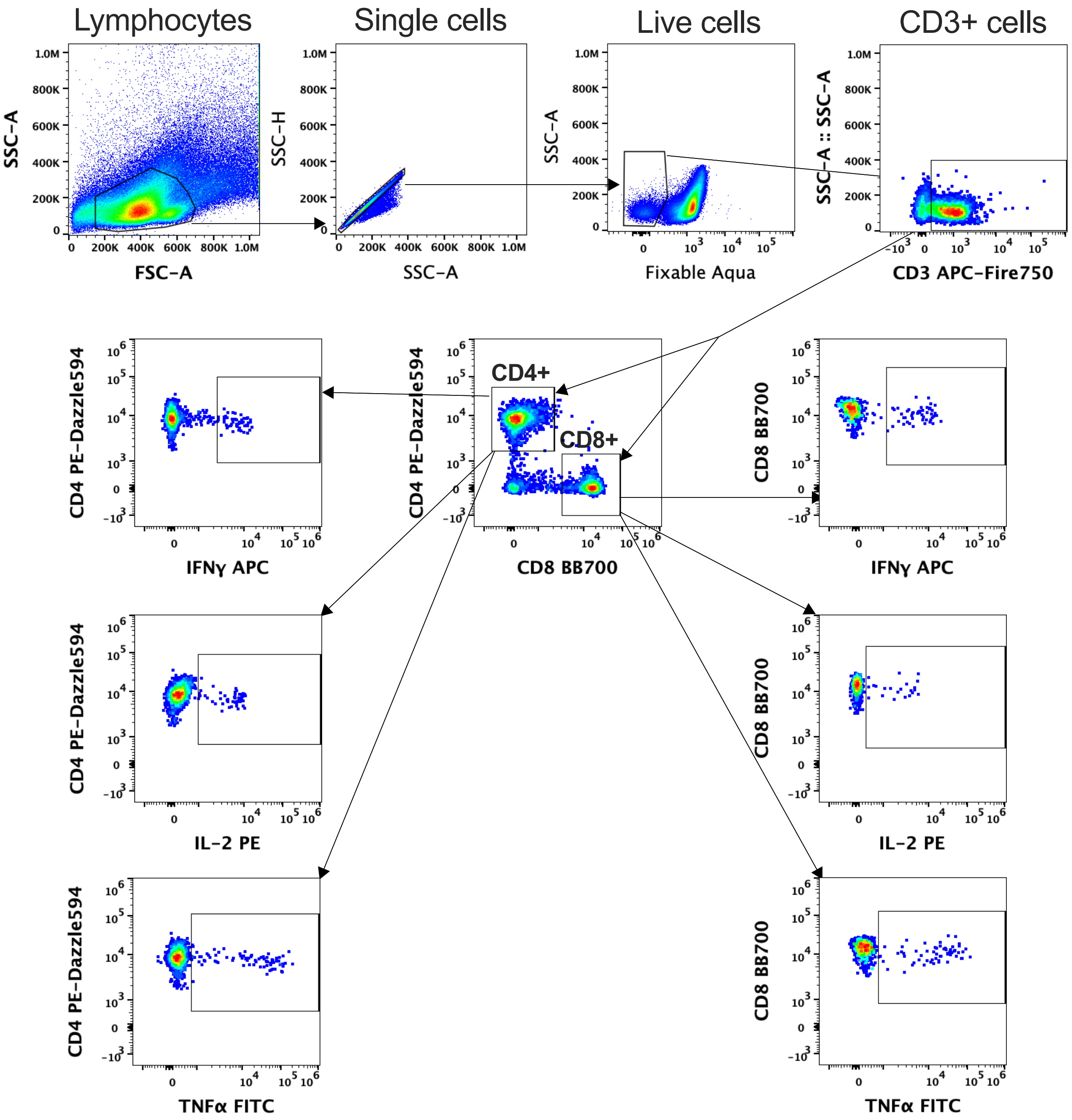


**Supplemental Figure 2 - Flow cytometry gating strategy**
Flow cytometry was performed on an Attune-NxT and data was analyzed with FlowJo version 10.10. The gating strategy for all samples is shown. Lymphocytes were gated on forward and side scatter area (FSC-A and SSC-A, respectively). Single lymphocytes were then gated on SSC area (SSC-A) and height (SSC-H). Dead cells, CD14+ monocytes and CD19+ B cells were then removed by fixable aqua, CD14 and CD19 staining. CD3+ T-cells were then gated on CD3 APC-Fire and SSC-A. CD4+ and CD8+ cells were separated on CD4 PE-Dazzle594 and CD8 BB700 staining. Within the CD4+ and CD8+ populations IFNγ, IL-2 and TNFα expression was assessed with IFNγ APC, IL-2 PE and TNFα FITC gating, respectively.

### Supplemental Figure 3

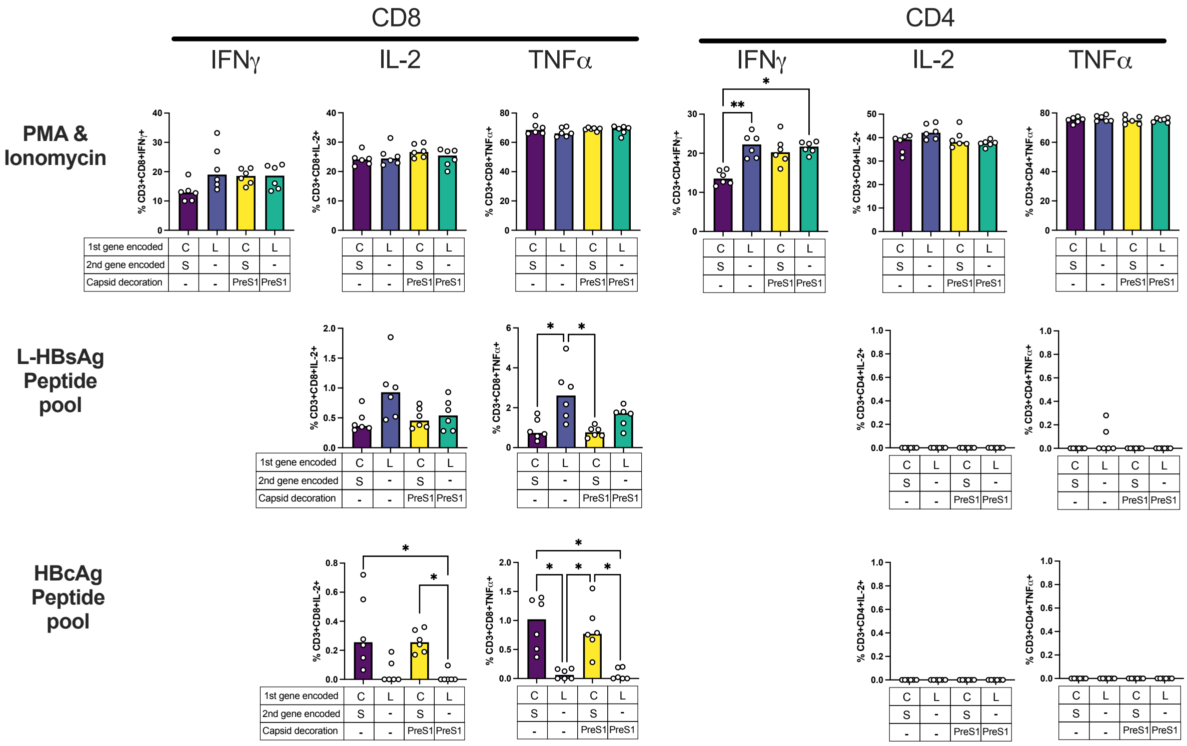


**Supplemental Figure 3 - Cytokine responses from CD4+ and CD8+ T-cells from the experiment shown in Figure 4**

Percentages of cytokine producing (IFNγ, IL-2 and TNFα) CD3+ CD4+ and CD3+ CD8+ T-cells at D35 after stimulation with either phorbol 12-myristate 13-acetate (PMA) and Ionomycin or peptide pools of L-HBsAg or HBcAg. Median responses are shown. Statistical analyses performed by Kruskal-Wallis with Dunn's test for multiple comparisons, **p<0.01, *P<0.05, non-significant comparisons are not shown. For HBcAg peptide pools, comparisons were only carried out between groups encoding the relevant antigens.

### Supplemental Figure 4

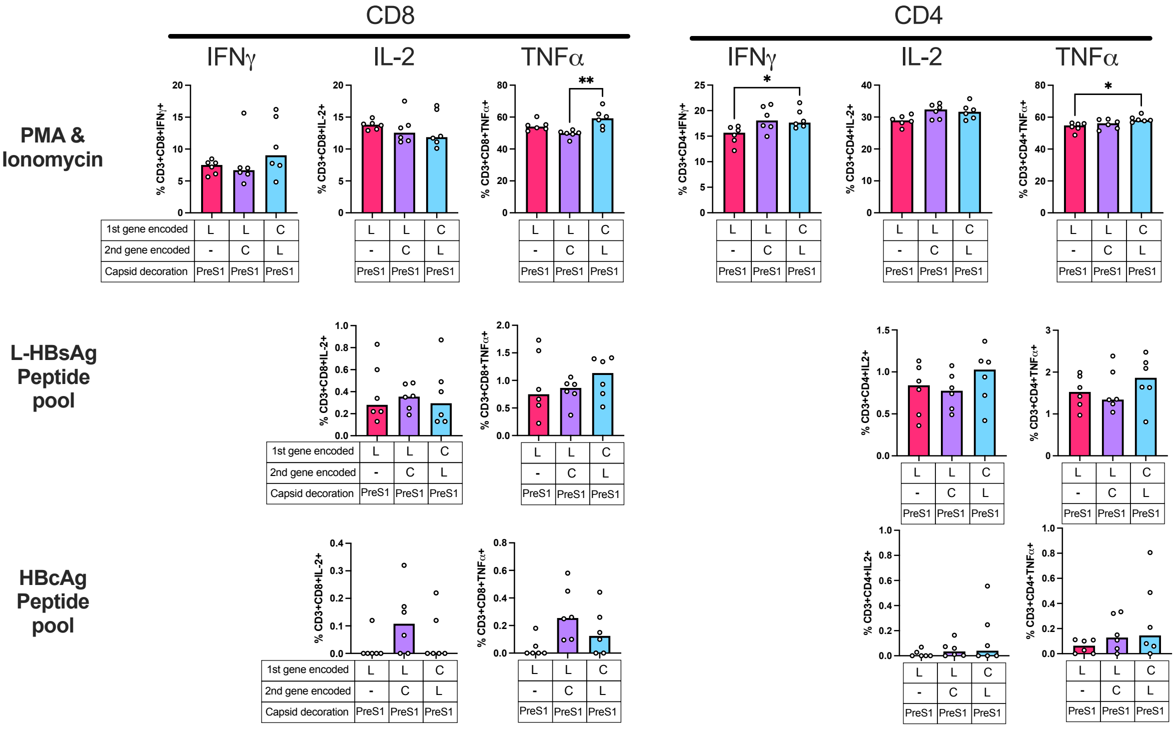


**Supplemental Figure 4 - Cytokine responses from CD4+ and CD8+ T-cells from the experiment shown in Figure 5**

Percentages of cytokine producing (IFNγ, IL-2 and TNFα) CD3+ CD4+ and CD3+ CD8+ T-cells at D35 after stimulation with either phorbol 12-myristate 13-acetate (PMA) and Ionomycin or peptide pools of L-HBsAg or HBcAg. Median responses are shown. Statistical analyses performed by Kruskal-Wallis with Dunn's test for multiple comparisons, **p<0.01, *P<0.05, non-significant comparisons are not shown. For HBcAg peptide pools, comparisons were only carried out between groups encoding HBcAg.
